# Effects of nicotine on the thermodynamics of the DPPC phase coexistence region

**DOI:** 10.1101/689588

**Authors:** Ernanni D. Vieira, A. J. Costa-Filho, Luis. G. M. Basso

**Author notes:** Corresponding author. E-mail address (Luis G. M. Basso).

## Abstract

Phase separation plays critical roles in several membrane functions, and reduction or disappearance of phase coexistence by action of membrane-interacting molecules have been implicated in membrane function impairment. Here, we applied differential scanning calorimetry, electron paramagnetic resonance (EPR), and non-linear least-squares (NLLS) spectral simulations to study the effects of nicotine, a parasympathomimetic drug, on the two-phase coexistence of dipalmitoyl phosphatidylcholine (DPPC) lipid membrane. The thermodynamic quantities describing the DPPC phase coexistence are temperature dependent, giving rise to non-linear van’t Hoff behavior. Our results showed that nicotine preferentially binds to the fluid phase and modifies the enthalpy and entropy changes of the DPPC heat capacity profile, while marginally perturbing the homogeneous gel and fluid phases. An EPR/NLLS/van’t Hoff analysis of the DPPC phase coexistence revealed that nicotine significantly modified the temperature dependence of the free energy change of the two-phase equilibrium from a cubic to a parabolic behavior, resulting in an alteration of the thermodynamical driving force and the balance of the non-covalent interactions of the lipids in equilibrium. The thermotropic behavior of the enthalpy, entropy, and heat capacity changes, as determined by EPR, indicated that nicotine modified the relative contributions of hydrogen-bonding, electrostatic interactions, and conformational entropy of the lipids to the thermodynamics of the phase coexistence. The predominantly entropically-driven gel-fluid transition in nicotine-free DPPC changes to a temperature-triggered entropically-driven or enthalpically-driven process in nicotine-bound DPPC. Further applications of this thermodynamic EPR/NLLS/van’t Hoff analysis are discussed.

## 1. Introduction

Biological membranes consist of complex mixtures of proteins, carbohydrates, and a wide variety of lipids with different hydrocarbon chains, polar groups, backbone structures, type of chemical linkages, etc [1]. Because of their complexity, studying biological membranes remains a major challenge from both experimental and theoretical perspectives. Therefore, a common strategy has been to investigate simple model membranes comprised of only one lipid or mixtures of lipids with the purpose of understanding the puzzle of membrane heterogeneity and intermolecular interactions. Calorimetric techniques such as differential thermal analysis (DTA) and differential scanning calorimetry (DSC) have been frequently used to investigate the physico-chemical properties of model and biological membranes [2]. Both techniques can provide direct information on the thermodynamics of the systems under study and, indirectly, on the molecular interactions in those systems [2]. DTA and DSC have also been used to study the effects of different additives such as drugs, polymers, peptides, and proteins in the lipid thermal transitions and in the corresponding phase diagrams of model and biological membranes [2−4]. However, the measure of heat flow does not allow for a complete description of the thermotropic processes at a molecular level. Therefore, specific structural alterations related to inter-and/or intramolecular interactions should be addressed using techniques other than DSC or DTA.

Electron paramagnetic resonance (EPR) of spin labels relies on the use of paramagnetic probes, mainly constituted by stable nitroxide radicals, covalently bound to the molecule of interest, and thus acting as "reporter" groups [5]. Spin labeled phospholipids have been widely used to monitor changes in membrane fluidity, packing of the hydrocarbon chains, local polarity surrounding the spin label, and phase transition temperatures, among other parameters, in model and biological membranes [6−9]. The sensitivity makes EPR one of the methods of choice when one intends to probe the thermotropic behavior of important microscopic parameters such as the local ordering and mobility of phospholipids in lipid bilayer membranes. The ability of EPR to track down alterations in the membrane dynamical structure can be further improved when simulations of the EPR spectra are included in the analyses [10-12]. Particularly, non-linear least-squares (NLLS) simulations are a powerful tool in decomposing multicomponent EPR spectra from spin probes partitioned into different phases that coexist in model and biological membranes [13‒16]. If the partition coefficient of the probe is known, the equilibrium constant *K* between the lipid states of a two-phase membrane is calculated by determining the fraction of the spin-labeled lipids in both states directly from NLLS spectral simulations. Therefore, van’t Hoff analyses can be performed upon recording changes in the equilibrium constant triggered by an external stimulus such as temperature. By plotting ln *K* versus reciprocal temperature 1/*T*, one can thus spectroscopically obtain important thermodynamic quantities such as the enthalpy change (Δ*H*^*0*^), the entropy change (Δ*S*^*0*^), the free energy change (Δ*G*^*0*^), and the heat capacity change (Δ*C*) of the process under investigation, without the need to directly measure heat flows. Therefore, EPR/NLLS in combination with van’t Hoff analysis emerge as an effective tool to indirectly extract meaningful thermodynamical information from the coexistence of phases displaying different EPR observables such as motional and/or structural properties in model and biological membranes.

We have previously applied this EPR/NLLS approach to study the thermodynamics of the ripple phase of pure dipalmitoyl phosphatidylcholine (DPPC) membranes and the phase coexistence region of a ternary mixture comprised of DPPC, palmitoyl–oleoyl phosphatidylcholine (POPC), and palmitoyl–oleoyl phosphatidylglycerol (POPG), which mimics the pulmonary surfactant lipid matrix [6]. Van’t Hoff analyses yielded non-linear curves, i.e., the dependence of ln *K* on 1/*T* was not linear over the temperature range corresponding to the phase coexistence region of the membranes, implying a temperature dependence of the thermodynamic parameters. Non-linear van’t Hoff plots have also been observed in several different biological applications [17−22].

In the present work, we seek to extend this method to study how small molecules such as drugs alter the thermodynamics, or ultimately, the van’t Hoff behavior of phase coexistence regions of model membranes. In particular, the drug of choice is nicotine, a natural product found in the nightshade family of plants, which is a potent parasympathomimetic stimulant acting as an agonist at most nicotinic acetylcholine receptors [23]. Although the mechanisms governing pulmonary nicotine absorption have been well studied [24], there is still no research on the effect of nicotine on the non-covalent interactions that stabilize the first barrier of protection of the pulmonary alveoli. The pulmonary surfactant matrix comprises of about 80% of phospholipids by weight, from which DPPC corresponds to approximately 50–70% [25]. Additionally, in eukaryotic cell membranes, phosphatidylcholine (PC) and phosphatidylethanolamine (PE) are the major lipid components, responsible for ca. 50 up to 70% of the lipid matrix in most membranes [26], with DPPC being one of the most abundant and studied PC lipids. Therefore, the lipid of choice in the present work was DPPC given its biological importance. In such a simple model membrane, the effects of foreign molecules, such as drugs [27, 28], peptides [29, 30] and proteins [31, 32] on the physico-chemical properties of the membrane provided a better understanding of the molecular details underlying those interactions.

Given that DPPC is the most abundant lipid in mammalian lung surfactant, we are particularly interested in investigating the effects of nicotine on the structural dynamics and on the thermodynamics of the two-phase region of DPPC bilayers, thus putatively mimicking its effect on the pulmonary lung surfactant matrix, which displays as main biological function the reduction of the surface tension at the air-liquid [33]. To do so, we used DSC and EPR spin label spectroscopy along with NLLS spectral simulations to investigate the thermotropic behavior of DPPC membranes in the presence of nicotine. Our experimental results show that nicotine profoundly alters the van’t Hoff behavior of the phase coexistence region of DPPC membranes and consequently the thermodynamic balance between the non-covalent interactions of the DPPC bilayer.

## 2. Materials and methods

### 2.1. Reagents

The phospholipid 1,2-dipalmitoyl-sn-glycero-3-phosphocholine (DPPC) and the spin label 1-palmitoyl-2-stearoyl(16-doxyl)-sn-glycero-3-phosphocholine (16-PCSL) were purchased from Avanti Polar Lipids, Inc. (Alabaster, AL). The (-)-1-methyl-2-(3-pyridyl)pyrrolidine was purchased from Sigma-Aldrich Brasil Ltda (Sao Paulo, Brazil). All reagents were used without further purification.

### 2.2. Sample preparation

Multilamellar vesicles of DPPC for DSC and EPR were prepared as described elsewhere [6]. Briefly, a measured volume of the nicotine stock solution in ethanol was added to DPPC (for DSC) or DPPC/16-PCSL (for EPR) in chloroform at different lipid:drug molar ratios: 50:1, 25:1, 10:1, 5:1, or 2:1. The organic solvent was vortexed thoroughly, evaporated under a N_2_ flow, and the resulting lipid film was placed under vacuum for 12 h. Pure lipid and lipid/nicotine samples were hydrated with a buffer containing 10 mM acetate, borate, phosphate, pH 7.4, sonicated for 10 min at 45 °C in a bath-type sonicator, and freeze-thaw cycled ten times prior to the experiments.

### 2.3. Differential scanning calorimetry (DSC) experiments

DSC measurements were carried out in a VP-DSC MicroCal microcalorimeter (Microcal, Northampton, MA, USA). The thermograms were recorded from 20 to 55 °C using a heating rate of 18.1 °C/h, after a 10-min equilibration period at 20 °C. The concentration of DPPC used was 2 mg/ml (2.72 mM). Data analysis were performed using Microcal Origin software.

### 2.4. Electron paramagnetic resonance (EPR) measurements

EPR experiments were carried out in a Varian E109 spectrometer operating at X-band (9.4 GHz) using the following acquisition conditions: microwave power, 10 mW; field modulation frequency, 100 kHz; field modulation amplitude, 0.2-0.5 G depending on the temperature range; sweep width, 160 G. The temperature was controlled using an E257-X Varian temperature control unit coupled to the spectrometer (uncertainty about 0.2 °C). The samples were transferred to glass capillaries (1.5 mm I.D.) and centrifuged at 10,000 rpm for 10 min. The capillaries containing the lipid pellets were then bathed in mineral oil (for thermal stability) inside an EPR quartz tube and placed in the center of the resonant cavity. Prior to measuring the EPR spectra, the sample was thermalized at each temperature for about three minutes. Typically, lipid samples contained 0.5 mg of DPPC and 0.5 mol% of 16-PCSL relative to the lipids.

### 2.5. Non–linear least–squares (NLLS) EPR spectral simulations

The NLSL software package, developed by Freed and co-workers [10], was used for the simulations of the EPR spectra of the 16-PCSL incorporated in the DPPC lipid bilayers. The most commonly used set of parameters in the calculation of the EPR spectrum are: the magnetic parameters, constituted by the *g*-tensor (*g*_*xx*_, *g*_*yy*_, *g*_*zz*_) and the hyperfine splittings (*A*_*xx*_, *A*_*yy*_, *A*_*zz*_), the components of the rotational diffusion tensor (*R*_*xx*_, *R*_*yy*_, *R*_*zz*_), and the ordering potential *U*(*Ω*), which is written as a series of spherical harmonic functions represented by the coefficients *c*_*20*_, *c*_*22*_, *c*_*40*_, *c*_*42*_, and *c*_*44*_[10,11]. In addition, in experiments with multilamellar vesicles the nitroxide-labeled lipid is considered microscopically ordered but macroscopically disordered and, therefore, the so-called microscopic order macroscopic disorder (MOMD) model is used [10,11]. The rotational diffusion tensor (*R*) of the spin-labeled lipid is described in terms of the local director of the bilayer, that is, it is calculated around axes parallel (*R*_*pll*_) and perpendicular (*R*_*prp*_) to the symmetry axis of the lipid acyl chain and the ordering potential, which describes the orienting influence of anisotropic fluids, such as membranes. As reported elsewhere [34], the 16-PCSL incorporated in lipid bilayers generally presents *R*_*pll*_≫ *R*_*prp*_, and, following a standard procedure [10, 12], we set *R*_*pll*_ = 10 *R*_*prp*_.

## 3. Results and Discussion

Membrane lipids can exist in a variety of organized states when hydrated [35]. Among the many factors that influence membrane polymorphism, temperature has been the most studied. DSC is one of the most extensively used techniques to study the thermotropic phase behavior and phase diagrams of biological model membranes [35]. The DSC thermogram exbhits the temperature dependence of the excess heat capacity of the membranes. The lipid acyl chain melting transition of one-component membranes is assumed to represent a simple two-state endothermic process, from which the transition midpoint temperature (*T*_*m*_), where the transition is 50% complete, can be directly measured and the calorimetric enthalpy change of the transition (Δ*H*_*cal*_) calculated from the area of the peak. At the *T*_*m*_, the change in the Gibbs free energy of the system is zero [36], allowing for the calculation of the entropy change (ΔS) associated with the transition.

Figure 1A shows representative DSC thermograms illustrating the thermotropic phase behavior of fully hydrated multilamellar DPPC vesicles in the absence and presence of nicotine. Pure DPPC displays two endothermic transitions, a low-enthapic and broad pretransition centered at *T*_*p*_ = 34 °C, and a very intense and more cooperative main phase transition, centered at *T*_*m*_ = 41 °C. The low-temperature peak arises from the transition of the solid−ordered lamellar gel phase, *L*_*β′*_, to the ripple gel phase, *P_β′_*, whereas the high-temperature peak arises from the conversion of *P_β′_* to the fluid, liquid crystalline *L*_*α*_ phase. In the *L*_*β′*_ phase, the lipids adopt an all-*trans* configuration and display a tilt angle of about 30° with respect to the membrane normal [37], whereas in the *L*_*α*_ phase, the lipid chains are mostly disordered and display many *trans* and *gauche* isomerizations in their C−C bonds. The *P*_*β′*_ phase is characterized by periodic undulations (ripples) on the membrane surface and is likely formed by periodic arrangements of both *L*_*β′*_ and *L*_*α*_ lipid domains [37]. The transition temperatures agree well with our previous works [6, 29].

**Figure 1.**
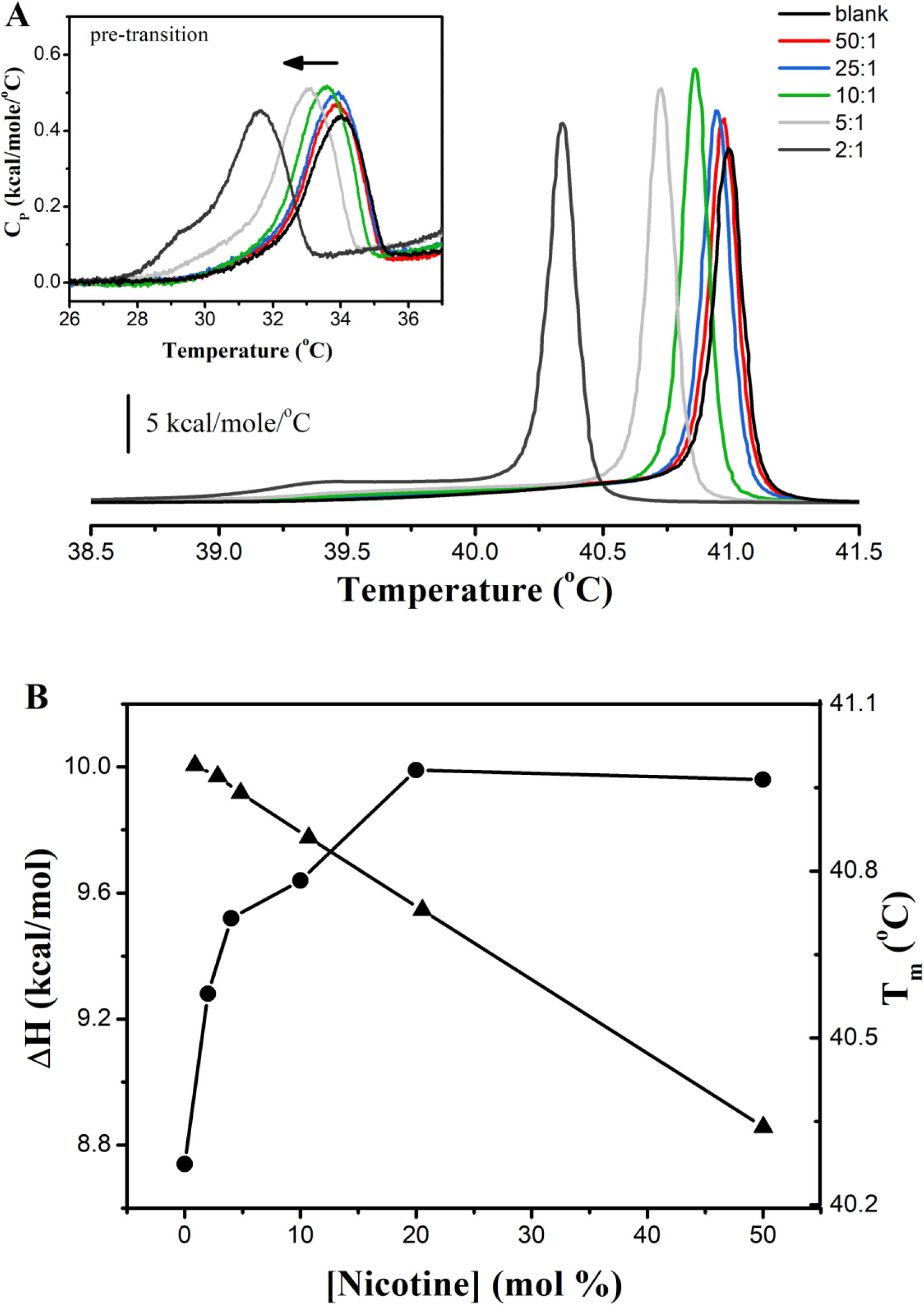
Thermotropic phase behavior of the membranes as monitored by DSC. (**A**) Temperature dependence of the molar heat capacity of DPPC in the absence and presence of nicotine at different lipid-to-drug molar ratios. The inset illustrates the pretransition of DPPC. (**B**) Effect of nicotine on the enthalpy change (Δ*H*, circles) and the melting temperature (*T*_*m*_, triangles) of DPPC lipid vesicles.

Addition of nicotine perturbs the heat capacity profile of DPPC in a concentration-dependent manner (Figure 1B). The thermograms of the nicotine-containing DPPC are progressively shifted towards low temperature, suggesting the drug preferentially distributes into the *L*_*α*_ phase of the membrane. Similar results were obtained with the antimalarial drug primaquine [27], whereas trans-parinaric acid caused an opposite effect, indicating a better partition into the gel phase [38]. Nicotine induced slight changes of the chain-melting temperature of DPPC/nicotine membranes (Δ*T*_*m*_< 1 °C), even at the highest lipid-to-drug molar ratio (2:1). This result implies that nicotine does not perturb the bilayer packing in a great extent. Indeed, the cooperativity of the transition, as qualitatively indicated by the inverse of the linewidth at half intensity (Δ*T*_*m,1/2*_), is virtually the same for all conditions (Table 1). Rigorously, the cooperative unit is calculated from the ratio of the van’t Hoff enthalpy change (Δ*H*_*vH*_) and Δ*H*_*cal*_. The average number of molecules in a cluster that melts cooperatively as a unit in the absence and presence of nicotine does not change considerably, ranging from 275 to 335 (Table 1).

**Table 1.**
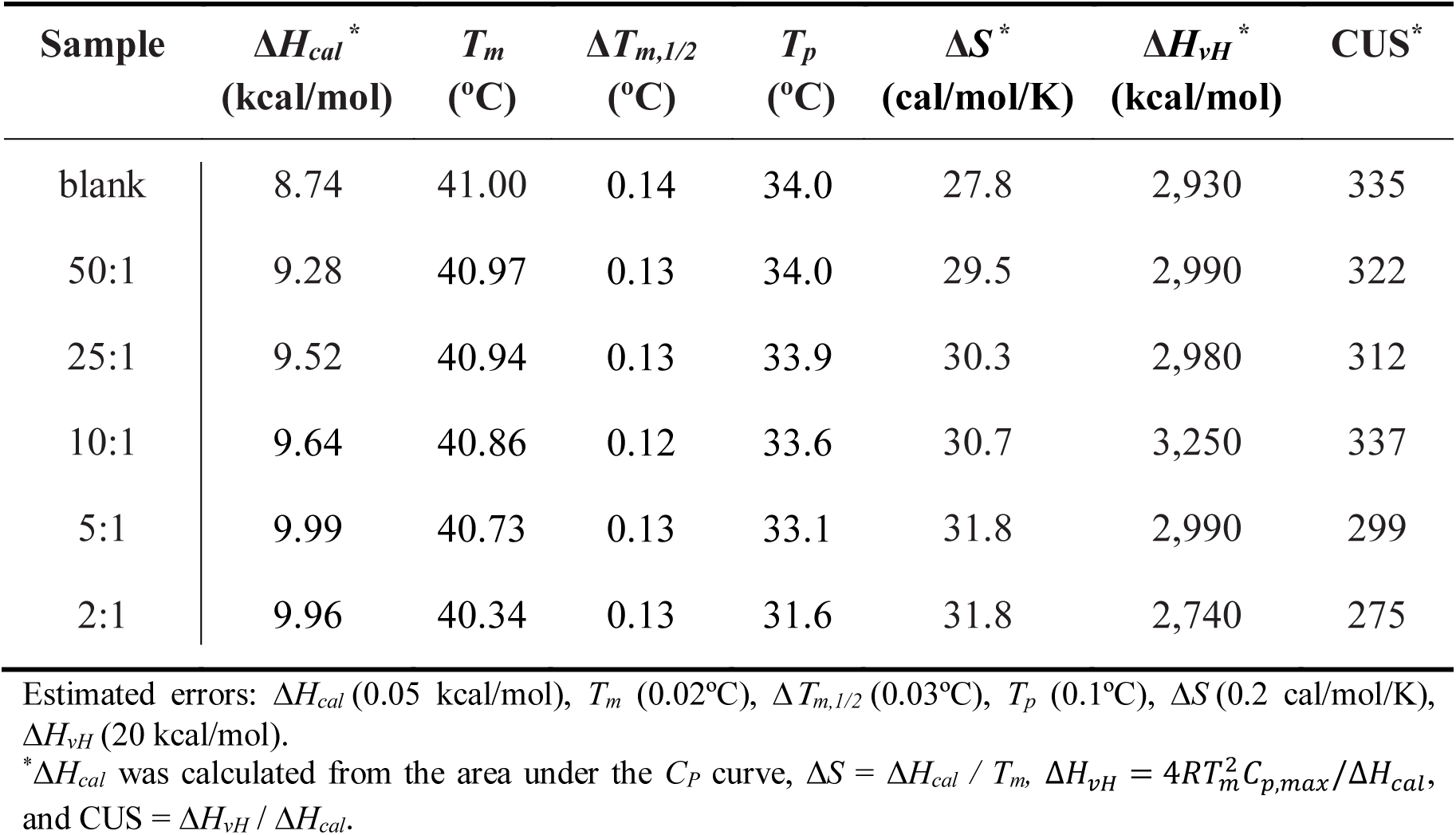
Calorimetric parameters obtained from the DSC thermogram of nicotine-free and nicotine-containing DPPC lipid vesicles at different lipid-to-drug molar ratios. Δ*H*_*cal*_ represents the enthalpy change of the whole phase transition curve; *T*_*m*_ is the main phase transition temperature; *Tp*, the pretransition temperature; Δ*T*_*m,1/2*_, the width at half height of the main phase transition peak; Δ*S* represents the entropy change of the transition at *T*_*m*_; Δ*H*_*vH*_, the van’t Hoff enthalpy; and CUS, the cooperative unit size, given in number of molecules.

| Sample | $\Delta H_{cal}^*$<br>(kcal/mol) | $T_m$<br>(°C) | $\Delta T_{m,1/2}$<br>(°C) | $T_p$<br>(°C) | $\Delta S^*$<br>(cal/mol/K) | $\Delta H_{vH}^*$<br>(kcal/mol) | CUS* |
| --- | --- | --- | --- | --- | --- | --- | --- |
| blank | 8.74 | 41.00 | 0.14 | 34.0 | 27.8 | 2,930 | 335 |
| 50:1 | 9.28 | 40.97 | 0.13 | 34.0 | 29.5 | 2,990 | 322 |
| 25:1 | 9.52 | 40.94 | 0.13 | 33.9 | 30.3 | 2,980 | 312 |
| 10:1 | 9.64 | 40.86 | 0.12 | 33.6 | 30.7 | 3,250 | 337 |
| 5:1 | 9.99 | 40.73 | 0.13 | 33.1 | 31.8 | 2,990 | 299 |
| 2:1 | 9.96 | 40.34 | 0.13 | 31.6 | 31.8 | 2,740 | 275 |

Estimated errors: $\Delta H_{cal}$ (0.05 kcal/mol), $T_m$ (0.02°C), $\Delta T_{m,1/2}$ (0.03°C), $T_p$ (0.1°C), $\Delta S$ (0.2 cal/mol/K), $\Delta H_{vH}$ (20 kcal/mol).
\* $\Delta H_{cal}$ was calculated from the area under the $C_p$ curve, $\Delta S = \Delta H_{cal} / T_m$ , $\Delta H_{vH} = 4RT_m^2 C_{p,max} / \Delta H_{cal}$ , and CUS = $\Delta H_{vH} / \Delta H_{cal}$ .

It has been shown that the pretrasition of phosphatidylcholines is very sensitive to the presence of foreign molecules at the membrane/water interface [27, 39]. Interestingly, binding of nicotine does not significantly alter either the *T*_*p*_ or the shape of the pretransition curve (Figure 1A, inset). This result suggests neither the structure of the DPPC head group [40] nor the membrane hydration [41, 42] are strongly affected by nicotine adsorption on the bilayer surface.

The most pronounced effects induced by nicotine were observed in the calorimetric enthalpy change of the whole transition, Δ*H*_*cal*_ (Figure 1B), and in the respective entropy change, Δ*S*, at the melting temperature *T*_*m*_(Table 1). Both Δ*H*_*cal*_and Δ*S* increase in a dose-dependent manner, but not exceeding 14% of the value obtained for the empty vesicles. These results indicate that more heat is needed to melt the acyl chains of nicotine-bound DPPC vesicles and more entropy is gained in this process.

Although accounting for a full thermodynamical description of the lipid phase transition, DSC does not provide the molecular details related to structural and dynamical changes taking place during the phase transition, which requires molecule-based methods to complement the DSC information.

EPR has been widely used to investigate the effect of temperature on model membranes [6,27-29,43,44]. Nitroxide-labeled phospholipids, such as n-PCSL (n=5, 7, 10, 12, 14, and 16, where n represents the position of the paramagnetic probe along the lipid acyl chain), are used to dissect the structural rearrangements experienced by the lipid matrix due to changes in the environment. Here, we chose the phospholipid 16-PCSL, spin-labeled at the 16^th^ C-atom of the sn-2 chain, because it has been shown to be very sensitive to detect phase coexistence between lipid micro-domains exhibiting different ordering and dynamics in EPR experiments [6,13]. Non-linear least-squares (NLLS) spectral simulations of the 16-PCSL EPR spectra were carried out to evaluate changes in the order parameter and the rotational diffusion rates of the lipid spin probe in the membrane. NLLS analysis is very effective in distinguishing the relative population of the spin probes between domains/phases with distinct membrane packing and fluidity [14−16, 27-29].

Figure S1 of the Supplementary Material shows the experimental and best-fit EPR spectra of 16-PCSL in hydrated multilamellar vesicles of DPPC in the absence and presence of nicotine at two lipid-to-drug molar ratios: 25:1 and 10:1. The EPR spectra recorded above ca. 40 °C from all samples displayed three narrow lines characteristic of a fast lipid probe in the disordered, liquid crystalline *L*_*α*_ phase of the bilayers. On the other hand, below a certain temperature that varied between 21 and 28 °C depending on the sample, the 16-PCSL EPR spectra exhibited broad lines, typical of spin probes in slow motion in the solid–ordered, *L*_*β′*_ phase of the DPPC membranes. In the temperature interval ranging from 28.3 to 40.1 °C for pure DPPC (Fig. S1A), 26.3 to 40.6 °C for the DPPC/nicotine 25:1 molar ratio (Fig. S1C) and from 21.1 to 38.8 °C for the DPPC/nicotine 10:1 molar ratio (Fig. S1E), the spectra presented overlap of broad and narrow components. A single spectral component could not be satisfactorily fitted to the EPR spectra recorded in those regions. The introduction of a second spectral population to the fitting process significantly improved the agreement between the experimental and theoretical curves (Figures S1B, S1D, and S1F). This result indicates phase polymorphism in those regions, that is, coexistence of micro-domains exhibiting different structural organization and membrane fluidity [6,27]. Indeed, it has been shown that the ripple phase of fully hydrated phosphatidylcholine lipid bilayers displays properties of both *L*_*β′*_ and *L*_*α*_ phases [45,46]. Interestingly, the appearance of the second spectral component takes place at a temperature lower than the onset temperature of the pretransition observed by DSC (about 29-30 °C, inset of Figure 1A), which may reflect changes in the 16-PCSL conformation due likely to microscopic lipid rearrangements of the surrounding lipids. This result shows the sensitivity of a molecule-based technique such as EPR in detecting different local structural changes before macroscopic endothermic events monitored by calorimetric techniques take place. The higher the nicotine concentration, the lower is the EPR-detected onset temperature and, thus, the larger is the temperature range corresponding to the phase coexistence region as compared to pure DPPC.

The order parameter (*S*) and the rotational diffusion rate (*R*_*prp*_), derived from the NLLS simulations, can provide valuable information on the structural arrangements of the bilayer and on the membrane fluidity, respectively. Figure 2 displays the temperature dependence of both parameters simulated with one and two spectral components. In general, the main phase transition is accompanied by abrupt changes in the order parameter (Figure 2A) and the acyl chain mobility (Figure 2B), whereas the pretransition exhibits mostly smooth structural changes of the lipid bilayer as seen by the paramagnetic end-chain lipid probe. Apart from the increased temperature range of the two-domain coexistence region, the EPR data also show that nicotine has little influence on the main phase transition temperature, confirming our DSC studies. Besides, the drug neither affects the ordering of the bilayer in the gel and fluid phases (Figure 2A) nor the fluidity of the gel phase (Figure 2B). The mobility of the lipids is only changed in the fluid phase at the highest lipid-to-drug molar ratio (10:1). Indeed, *R*_*prp*_ increases by 35-40% in the fluid phase of DPPC/nicotine membranes as compared to the same phase in the nicotine-free vesicles. This result suggests that nicotine may act as a lipid spacer when adsorbed at high concentrations onto the DPPC surface in the liquid crystalline phase, thus increasing the membrane fluidity of the lipid bilayer. However, nicotine does not alter the activation energy for the 16-PCSL in both gel and fluid phases (Table S1), suggesting that the drug has no effect on the sensitivity to the temperature of the reorientation motion of the molecular axis perpendicular to the long axis of the lipid spin probe in both phases [47]. This result implies that the microviscosity around the nitroxide in the bilayer midplane is virtually the same in the absence and presence of nicotine, yielding similar temperature dependence of the rate of mobility change.

**Figure 2.**
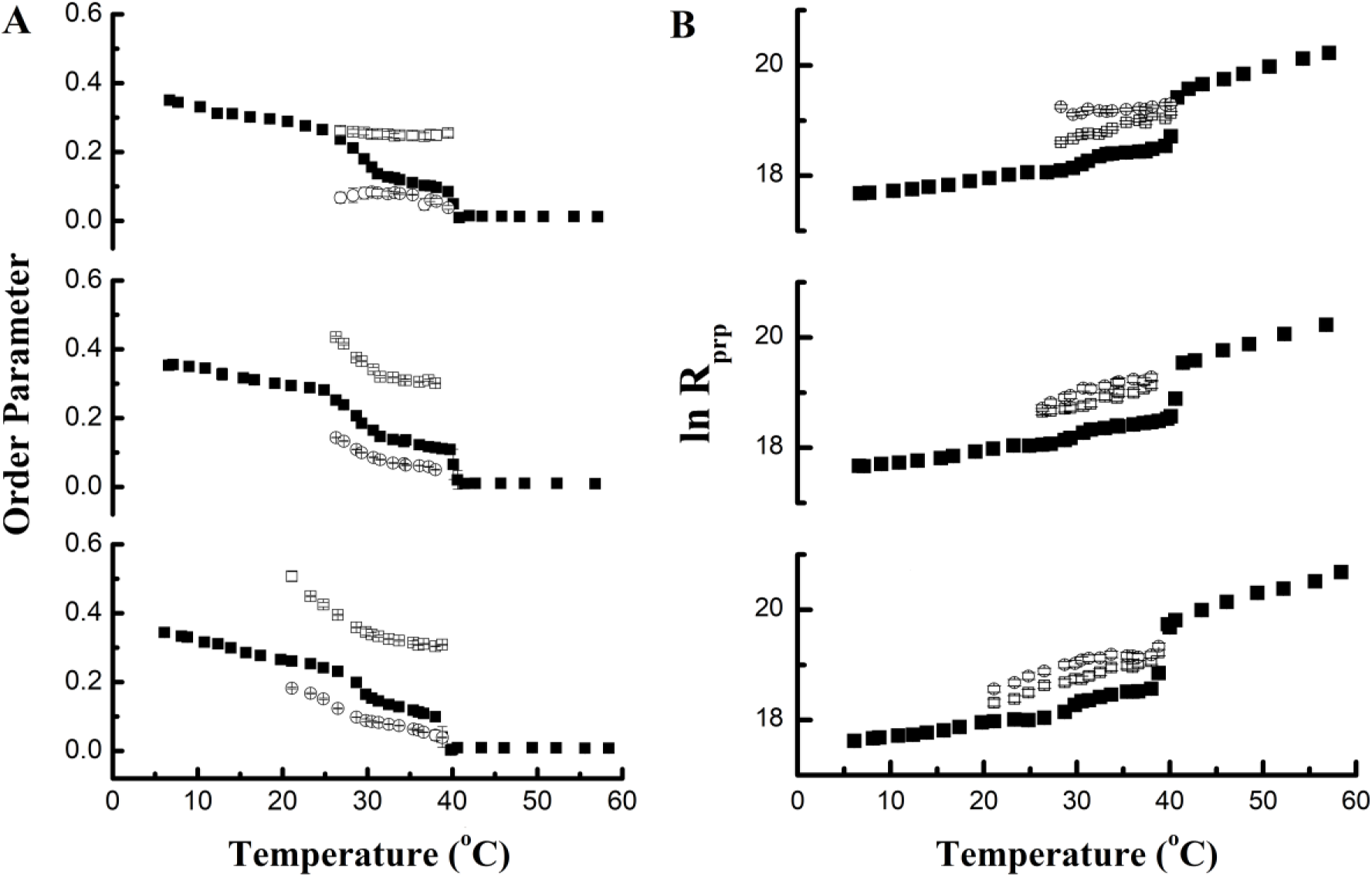
Thermotropic phase behavior of the membranes as monitored by EPR. Temperature dependence of **(A)** the order parameter *S* and **(B)** the rotational diffusion rate *R*_*prp*_ of 16-PCSL in pure DPPC (top panels) and DPPC/nicotine lipid vesicles at two lipid-to-drug molar ratios: 25:1 (middle panels) and 10:1 (bottom panels). Both parameters were obtained from the NLLS simulations of the EPR spectra using one (full black squares) or two (open squares and circles) spectral components.

In the phase coexistence region, the major effect of nicotine is a marked increase in the order parameter of the more ordered *L*_*β′*_ phase (Figure 2A). The order parameter of the gel-phase component in the empty DPPC membranes, which is virtually temperature invariant (S ~ 0.25−0.26) over the entire coexistence region, assumes much higher values (S ~ 0.3−0.5) in the presence of nicotine and becomes linearly dependent on temperature. This result indicates that nicotine induces a membrane structural reorganization by significantly enhancing the packing of the ordered domain in the ripple phase. On the other hand, the ordering of the fluid domain is only slightly affected by nicotine. We hypothesize that the more packed ordered domain along with the larger temperature range corresponding to the phase coexistence might be the molecular mechanisms behind the enthalpy and entropy gains during phase transition induced by nicotine as compared to the empty membranes, as shown by our DSC experiments. Indeed, more heat is needed to melt more lipids in the *L*_*β′*_ phase and additional entropy is gained in the conformational change between the more ordered, drug-induced, *trans*-rich lipids to the disordered, *gauche*-rich lipids.

We can gain further knowledge about the thermodynamics of the DPPC/nicotine system by applying a different theoretical analysis of the data corresponding to the two-domain coexistence region. The population of each spectral component is an important parameter to extract thermodynamical information of the nicotine/lipid system. They can be treated on the basis of a van’t Hoff analysis by measuring the equilibrium constant in the phase coexistence region at each temperature. Considering *K* the equilibrium constant between the lipids in the *L*_*β′*_ and *L*_*α*_ domains of the DPPC ripple phase and *K*_*P*_ the partition coefficient of 16-PCSL in both domains, then [6]

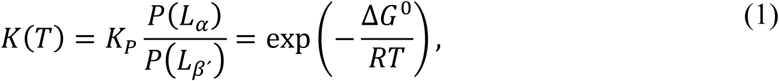

where *P*(*L*_*α*_) and *P*(*L*_*β′*_) represent the populations of 16-PCSL in the *L*_*α*_ and *L*_*β′*_ phases, respectively, obtained by NLLS spectral simulations, *R* is the universal gas constant, and Δ*G^0^* corresponds to the free energy change necessary to transfer a lipid from the gel state to the fluid state. A plot of ln *K* versus 1/*T* (van’t Hoff plot) can thus provide the thermodynamic quantities associated to the phase coexistence region. The van’t Hoff analysis has been classified as classical when the plot of ln *K* versus 1/*T* is linear and nonclassical otherwise [48,49]. Deviation from linearity has been observed in several systems [17−22], which means that Δ*G*^*0*^, the standard enthalpy (Δ*H*^*0*^), standard entropy (Δ*S*^*0*^), and standard heat capacity (Δ*C*^*0*^) changes are temperature dependent [48,49]. We have previously shown that the dependence of ln *K* on 1/*T* over the temperature range corresponding to the phase coexistence region of DPPC and a pulmonary surfactant lipid model membrane is non-linear and can be written as [6]:

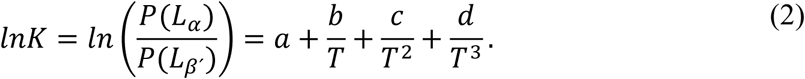

In the above equation, we assumed that 16-PCSL partitions equally well between the gel and fluid phases of DPPC and is temperature independent, i.e., *K*_*P*_= 1, as previously reported [15,50−52]. The empirical coefficients *a*, *b*, *c*, and *d* are estimated by fitting Eq. 2 to the experimental ln *K* versus 1/*T* curves and the thermal behavior of the thermodynamic quantities (Δ*G*^*0*^, Δ*H*^*0*^, Δ*S*^*0*^, and Δ*C*^*0*^) are determined as described elsewhere [6].

Figure 3 illustrates the van’t Hoff plots for the 16-PCSL embedded in the membranes studied. The temperature dependence of ln *K* obtained for the DPPC membranes was best fitted with a cubic function, as previously observed [6], whereas the effect of nicotine on the gel-fluid equilibrium yielded a parabolic van’t Hoff plot. This result is remarkable: qualitatively, binding of nicotine to DPPC membranes affects the van’t Hoff behavior of the DPPC/nicotine ripple phase in a way similar to that caused in the *L*_*α*_ −*L*_*β′*_ coexistence region of the ternary DPPC/POPC/POPG mixture at 4:3:1 molar ratio due to addition of the unsaturated POPC and POPG lipids to DPPC membranes [6]. Nicotine significantly changed the van’t Hoff plots at lower temperatures, indicating different thermal behavior of the spectral populations. Binding of nicotine induces the appearance of a higher population of fluid lipids (50-55%) as compared to the drug-free membranes (~ 30%) at the onset temperature where phase coexistence is resolved by EPR. Upon heating, the fluid population of the ripple phase of pure DPPC membranes increases in a non-linear fashion and tends to 50% at *T*_*m*_, whereas molecular rearrangements take place in the DPPC/nicotine ripple phase in such a way that the fluid population decreases to about 37-39% at *T*_*P*_ and then increases to 50% at *T*_*m*_.

**Figure 3.**
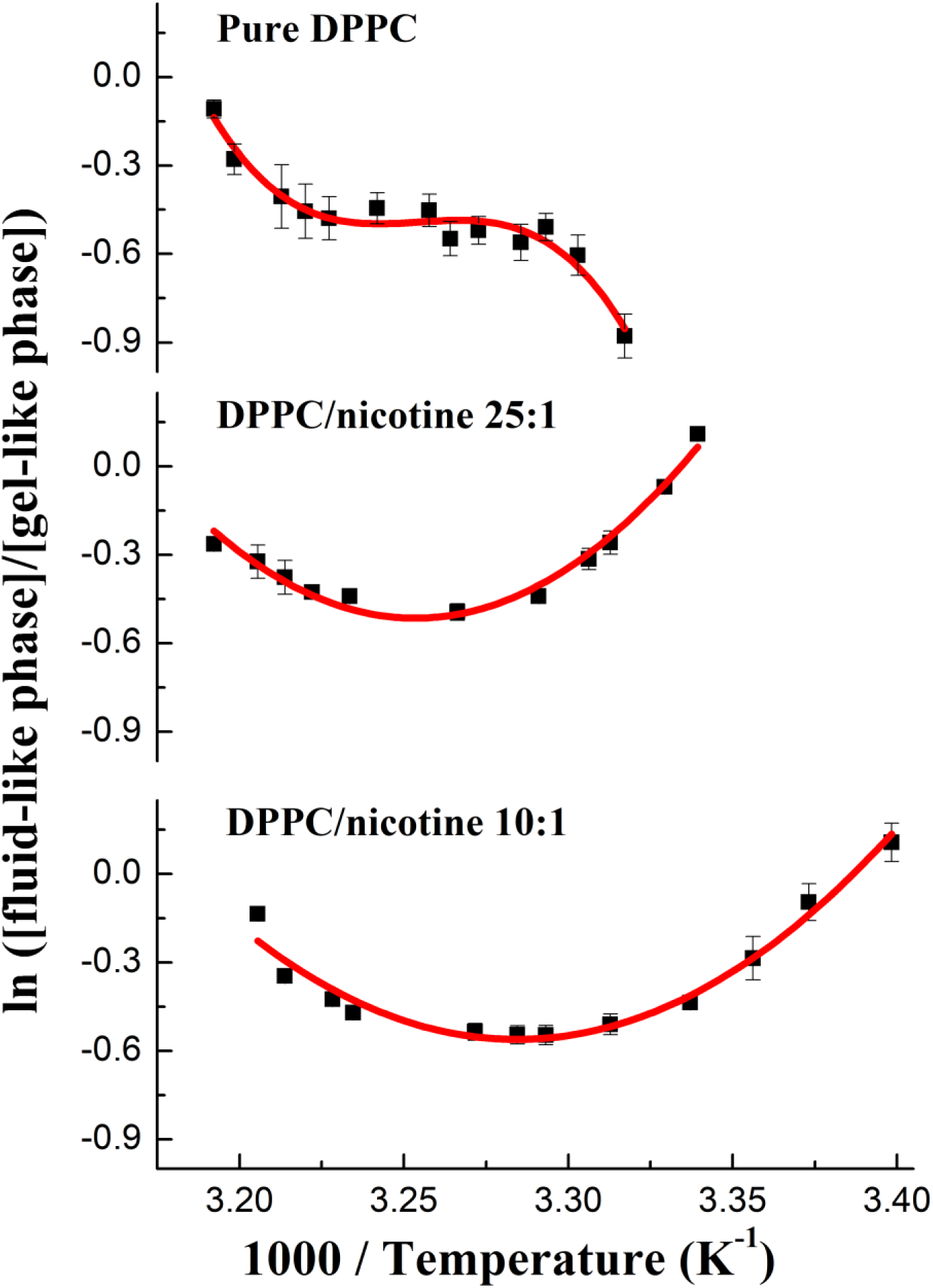
Van’t Hoff analysis of pure DPPC and DPPC/nicotine multilamellar vesicles. The solid lines are the best fits to the van’t Hoff plots. Pure DPPC was best fitted with a cubic function, whereas the effect of nicotine yielded a parabolic van’t Hoff plot. The lipid-to-drug molar ratios were 25:1 and 10:1.

The alteration in the non-linear van’t Hoff plots of DPPC in the presence of nicotine also leads to different thermotropic behaviors of Δ*G^0^*, Δ*H^0^*, Δ*S^0^*, and Δ*C^0^* associated to the equilibrium between the gel and fluid phases, as shown in Figures 4 and 5. The best-fit coefficients of the ln *K* versus 1/*T* curves (*a*, *b*, *c*, and *d;* Figure 3) are shown in the Table S2 of the supplementary material. As can be observed in Figures 4A and 5A, interconversion of gel-state lipids to fluid-state lipids is more thermodynamically favorable in the presence of nicotine for *T <* 32−33°C, i.e., Δ*G^0^* is lower in the presence of the alkaloid and further decreases upon cooling, whereas it increases with lowering *T* for pure DPPC. On the other hand, for *T* > 33−34°C, all Δ*G*^*0*^ curves display similar behavior with temperature and tend to zero as *T* approaches *T*_*m*_. Interestingly, the saddle point of the Δ*G*^*0*^ curve for the empty DPPC membranes (Figure 4A) and the maximum of the curves obtained in the presence of nicotine (Figure 5A) correspond to the pretransition temperature, *T*_*P*_, of the membrane systems and can be determined by taking the first derivative of Δ*G*^*0*^ relative to *T* and setting it to zero. Since (∂∆*G*^0^/∂*T*)_*p*_ = − ∆*S*^0^, the entropy change associated to the isobaric transfer of 16-PCSL from the solid-ordered gel phase to the disordered fluid phase is zero at *T*_*P*_. This result implies that the entropy gained in the conformational change from *trans* to *gauche* isomerizations and in the formation of the membrane undulation of the ripple phase is counterbalanced, at *T*_*P*_, by the interaction of the head group of the fluid lipids with solvent molecules, which contributes with a negative Δ*S*^*0*^.

**Figure 4.**
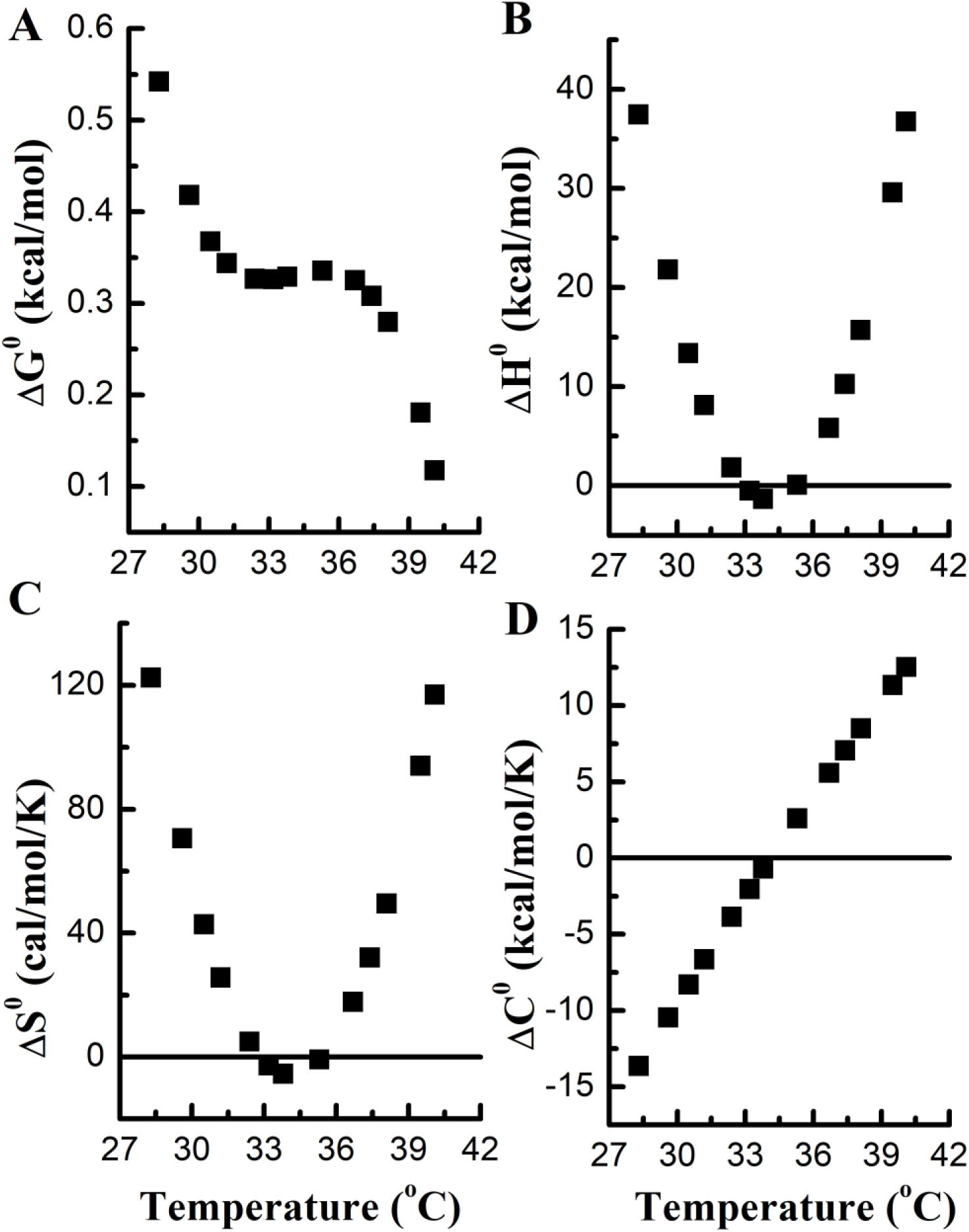
Thermodynamics of the phase coexistence region of DPPC vesicles. Temperature-dependence of the changes in the (**A**) Gibbs free energy, Δ*G*^*0*^, (**B**) enthalpy, Δ*H*^*0*^, (**C**) entropy, Δ*S*^*0*^, and (**D**) heat capacity, Δ*C*^*0*^.

**Figure 5.**
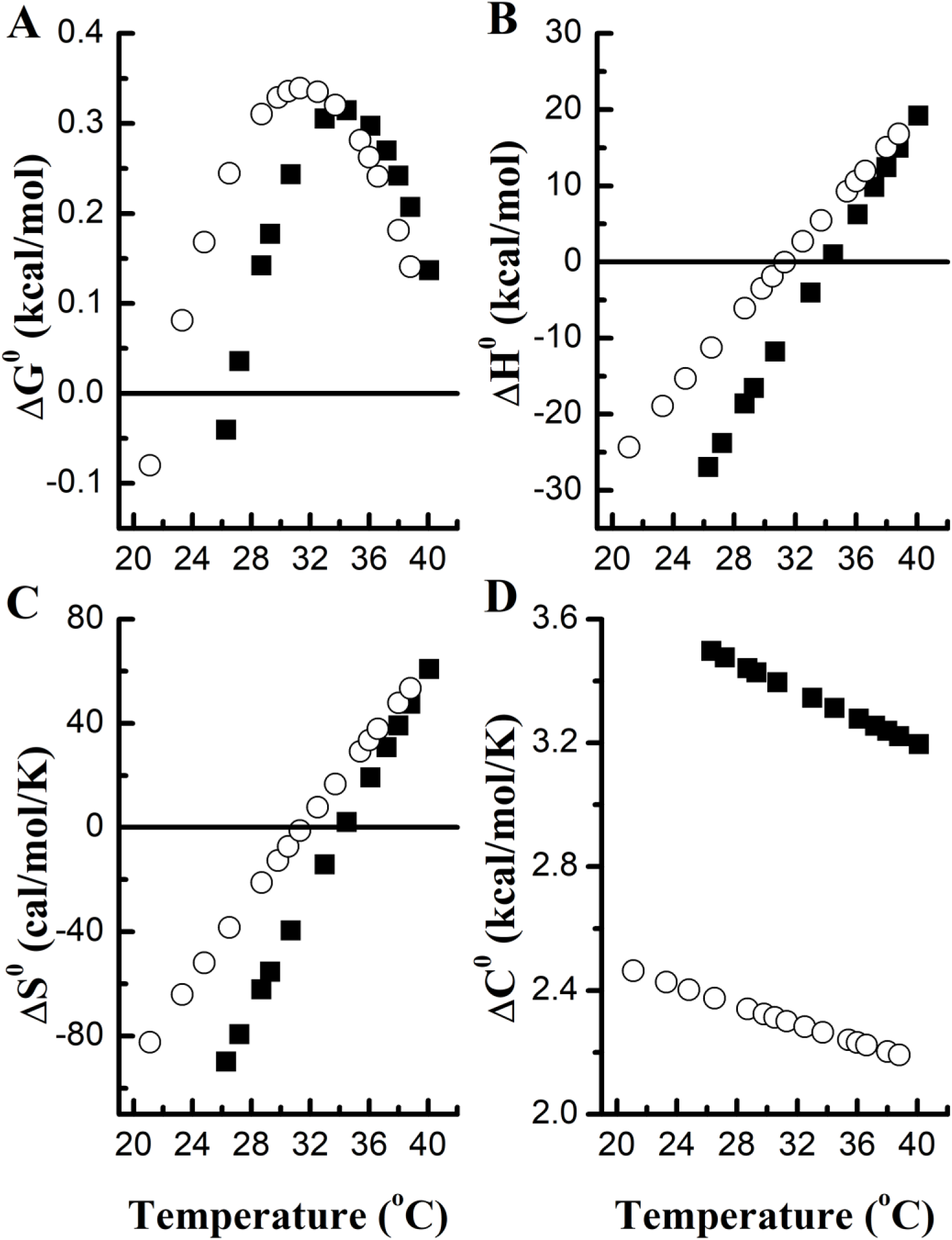
Thermodynamics of the phase coexistence region of DPPC/nicotine vesicles. Temperature-dependence of the changes in the (**A**) Gibbs free energy, Δ*G*^*0*^, (**B**) enthalpy, Δ*H*^*0*^, (**C**) entropy, Δ*S*^*0*^, and (**D**) heat capacity, Δ*C*^*0*^ for nicotine-containing membranes at lipid-to-drug molar ratios of 25:1 (solid squares) and 10:1 (open circles).

The enthalpy and entropy changes of the two-phase coexistence region are also dramatically modified by nicotine. The Δ*H^0^* and Δ*S^0^* curves obtained for pure DPPC change from a predominantly positive parabolic dependence with the temperature (Figures 4B and 4C) to a linear behavior with *T* in the presence of nicotine, varying from negative to positive values depending on the temperature (Figures 5B and 5C). These results are likely due to differences in the heat capacity changes between the gel and fluid domains in the absence and presence of the alkaloid, since ∆*C*^0^ = (∂∆*H*^0^/∂*T*)_*p*_ = *T* (∂∆*S*^0⁄^∂*T*)_*p*_. Δ*C*^*0*^ varies linearly from large negative values, for *T* < *T*_*P*_, to large positive values upon raising the temperature for the empty membranes (Figure 4D), whereas it is positive and slightly linearly dependent on *T* for the nicotine-embedded membranes in the phase coexistence region (Figure 5D). Additionally, Δ*C*^*0*^ is reduced with increased nicotine concentration. Taken together, these findings indicate that nicotine substantially modifies the thermodynamical driving force in the phase coexistence region of DPPC membranes. Indeed, while the thermotropic process associated with the gel−fluid equilibrium (for *T* < *T*_*m*_) in pure DPPC is predominantly entropically driven (Δ*H*^*0*^ > 0 and Δ*S*^*0*^ > 0), nicotine induces an entropically−driven process for *T* > *T*_*P*_ and an enthalpically−driven process for *T* < *T*_*P*_ (Figure 6). Despite that, entropy–enthalpy compensation still holds in the DPPC/nicotine membranes (Figure 6), which indicates that drug binding and membrane perturbation give rise to a very small contribution to the free energy changes between the gel and fluid states, as previously shown for membrane-interacting proteins [53].

**Figure 6.**
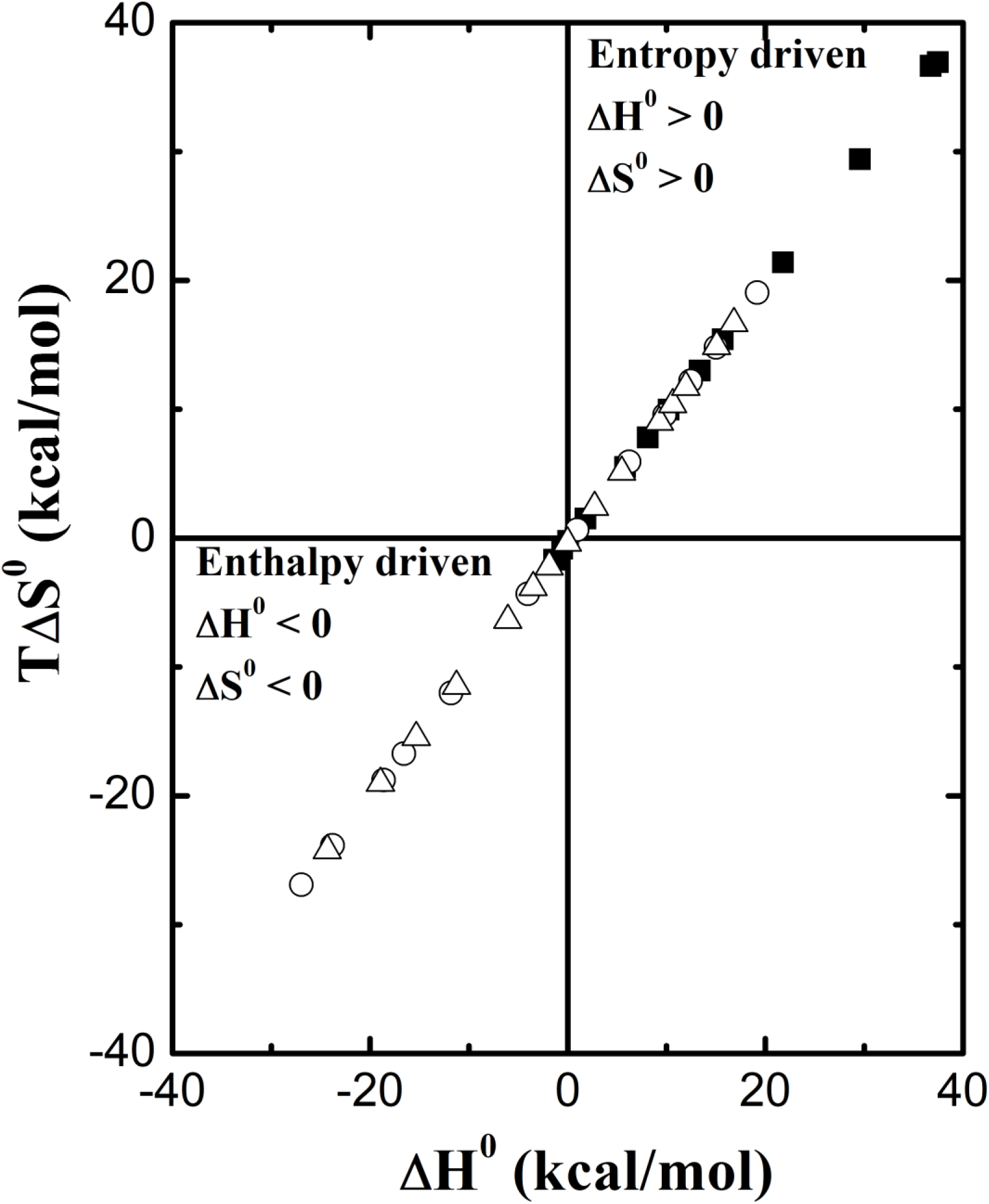
Entropy-enthalpy compensation of the DPPC/nicotine vesicles. *T*Δ*S*^*0*^ хΔ*H*^*0*^ plots for pure DPPC (solid squares) and DPPC/nicotine vesicles at lipid-to-drug molar ratios of 25:1 (open circles) and 10:1 (open triangles).

At physiological or mildly acidic pH values, nicotine is a monoprotonated cation with the positive charge centered on the pyrrolidine nitrogen atom [54]. Adsorption of nicotine onto the DPPC membranes should thus be restricted to the membrane/water interface and may likely occur through insertion of the pyridine ring in the bilayer in a wedge−like conformation, since the pyridine and pyrrolidine rings seem to be oriented roughly perpendicular to one another [55]. The alkaloid does not alter the structure of the DPPC head group and the average hydration property of the lipid bilayer, as demonstrated by the lack of prominent effects on the lipid pretransition (Figure 1A). However, since the positively charged pyrrolidinyl nitrogen may interact electrostatically with the negatively charged phosphate group and with the carboxylate ester groups within the lipid head group, nicotine may compete with water molecules at the membrane surface. Dislocation of water molecules from the lipids may take place but is likely counterbalanced by the nicotine-induced shift of the equilibrium constant towards the formation of fluid lipids, which bind more water molecules than lipids in the gel state [56]. The alkaloid thus changes the hydrogen bonding network in the polar head group region and may stabilize defects at the membrane−water interface [57]. These might be the interfacial contributions to the entropy and enthalpy changes between the gel and fluid lipids induced by nicotine. Moreover, the contribution from the acyl chains to both Δ*H*^*0*^ and Δ*S*^*0*^ of the phase coexistence region is two-fold: 1) negative, due to the substantial ordering effect caused by nicotine in the solid-ordered lipids in the *P*_*β′*_ phase; and 2) positive, due to the higher population of fluid lipids (increased *gauche*-to-*trans* conformer ratio) promoted by nicotine as compared to the empty membranes [58]. The ordering effect is likely triggered by electrostatic interactions in the polar region, thus enhancing the van der Waals interactions. As the temperature increases thermal fluctuation gradually decreases the order parameter of the more ordered component in the nicote-containing membranes and changes *K* so that it modulates the relative contributions of the head group and acyl chains to both Δ*H*^*0*^ and Δ*S*^*0*^ values.

Our findings show that the interaction of nicotine with the DPPC lipid bilayer changes the thermotropic behavior of the thermodynamic quantities associated with the two-phase coexistence region in a qualitatively similar way to what was observed in the studies of the ternary 4:3:1 DPPC/POPC/POPG mixture, which mimics the lipid bilayer matrix of the pulmonary surfactant complex [6]. Comparatively to our previous study [6], we found the following from the non-linear van’t Hoff analyses: 1) incorporation of nicotine onto pure DPPC substantially modifies the thermodynamic parameters of the two-phase region of the membrane in a way qualitatively similar to the addition unsaturated POPC and POPG lipids into pure DPPC; 2) increasing nicotine concentrations reduced the heat capacity change of the DPPC ripple phase in such a way that resembles the effect of increasing ionic strengths in the DPPC/POPC/POPG mixture; and 3) the positively charged alkaloid seems to have a similar effect in DPPC membranes as sodium cations in the ternary lipid mixture. Nicotine-induced changes in the van’t Hoff plots of DPPC led to marked alterations of the temperature dependence of Δ*G*^*0*^, Δ*H*^*0*^, Δ*S*^*0*^, and Δ*C*^*0*^ of the two-phase coexistence region likely due to modifications in the hydrogen-bonding properties and the conformational entropy of the lipids.

In the present work, we extended our previous EPR/van’t Hoff analysis of lipid model membranes exhibiting coexistence of *L*_*β’*_ −*L*_*α*_ phases [6] to incorporate the study of the interaction of a small, drug-like molecule with a two-phase membrane. Increasing evidence shows that the particular composition of biological membranes gives rise to a lateral segregation of their constituents into liquid-disordered and raft-like, liquid-ordered phases, with the latter being enriched in proteins, cholesterol, and sphingolipids and implicated in a number of important dynamic cellular processes [59]. In particular, the pulmonary surfactant matrix has been shown to display lateral phase separation up to physiological temperature [60]. Ordered/disordered phase coexistence is thought to be critical for the lung surfactant biophysical function [61]. The interaction of external agents such as drugs, amphiphilis, and/or proteins with surfactant membranes may not only alter the structure and fluidity of the membranes but also disturb the equilibrium between the ordered and disordered domains, potentially leading to complete disappearance of the phase coexistence. Indeed, the surfactant inhibitor C-reactive protein has been shown to bind to surfactant membranes, increase membrane fluidity, and abolish the ordered/disordered phase coexistence, resulting in lung surfactant inactivation [62]. Therefore, we believe that our thermodynamic EPR method might also be useful to shed light on the thermodynamics of complex biological membranes exhibiting lateral phase separation and to investigate how external agents alter the thermodynamical driving force of the phase equilibrium.

## 4. Conclusion

We employed DSC and EPR along with NLLS spectral simulations to study the interaction of nicotine with pure DPPC lipid bilayers. The drug preferentially binds to the fluid phase but does not change the melting profile of the membrane in a prominent way. On the contrary, nicotine slightly increases the enthalpy and entropy changes of the DPPC acyl chain melting transition. NLLS simulations of EPR spectra revelead that the structure, dynamics, and activation energies of the gel and fluid phases are marginally perturbed by the drug. On the other hand, nicotine profoundly alters the van’t Hoff behavior of DPPC in the gel-fluid coexistence region. The non-linear, cubic dependence of the free energy change with the temperature is changed to a parabolic behavior in the presence of nicotine. Van’t Hoff analysis showed that the thermodynamical driving force and the balance of the non-covalent interactions of the lipids in equilibrium in the ripple phase of the DPPC bilayer matrix are substantially altered. Transition from a gel state to a fluid state in pure DPPC is predominantly entropically-driven, while nicotine-lipid interactions yield both entropically- and enthalpically-driven processes depending on the temperature. The thermal behavior of the heat capacity change in the phase coexistence region of pure DPPC and DPPC/nicotine lipid bilayers suggested that interaction of nicotine with DPPC alters the relative contributions of the conformational entropy of the lipids and the electrostatic interactions and the H-bond formation in the lipid head group. We believe this thermodynamic EPR approach could be extended to study the interaction of membrane-active molecules with more complex model and biological membranes provided coexistence of different domains could be EPR detected.

## Acknowledgements

The authors acknowledge the Brazilian agencies FAPESP (Grant nos. 2010/17662-8, 2015/18390-5, 2015/50366-7), CNPq (Grant nos. 475696/2013-1, 308380/2013-4) and CAPES for financially supporting this work. LGMB thanks FAPESP for a postdoctoral fellowship (2014/00206-0). The authors are also grateful to the “Sergio Mascarenhas” Molecular Biophysics Group at Sao Carlos Physics Institute of the University of Sao Paulo for allowing access to the DSC and EPR facility.

