## Supplementary Material for "Effects of nicotine on the thermodynamics of the DPPC phase coexistence region"

[b]Laboratório de Biofísica Molecular, Departamento de Física, Faculdade de Filosofia, Ciências e Letras de Ribeirão Preto, Universidade de São Paulo, Ribeirão Preto, SP – Brasil

* Corresponding author. Phone./fax: +55 (16) 3373-8094.

**Figure S1.** Experimental (black) and best-fit (red) EPR spectra of 16-PCSL in pure DPPC (**A** and **B**) and in DPPC/nicotine lipid vesicles at two lipid-to-drug molar ratios: 25:1 (**C** and **D**) and 10:1 (**E** and **F**). Only representative spectra at selected temperatures are illustrated. The spectra in panels A, C and D were simulated with only one component, whereas those in panels B, D and F were fitted with two components (green and blue). Arrows point to the second component in the phase coexistence region.

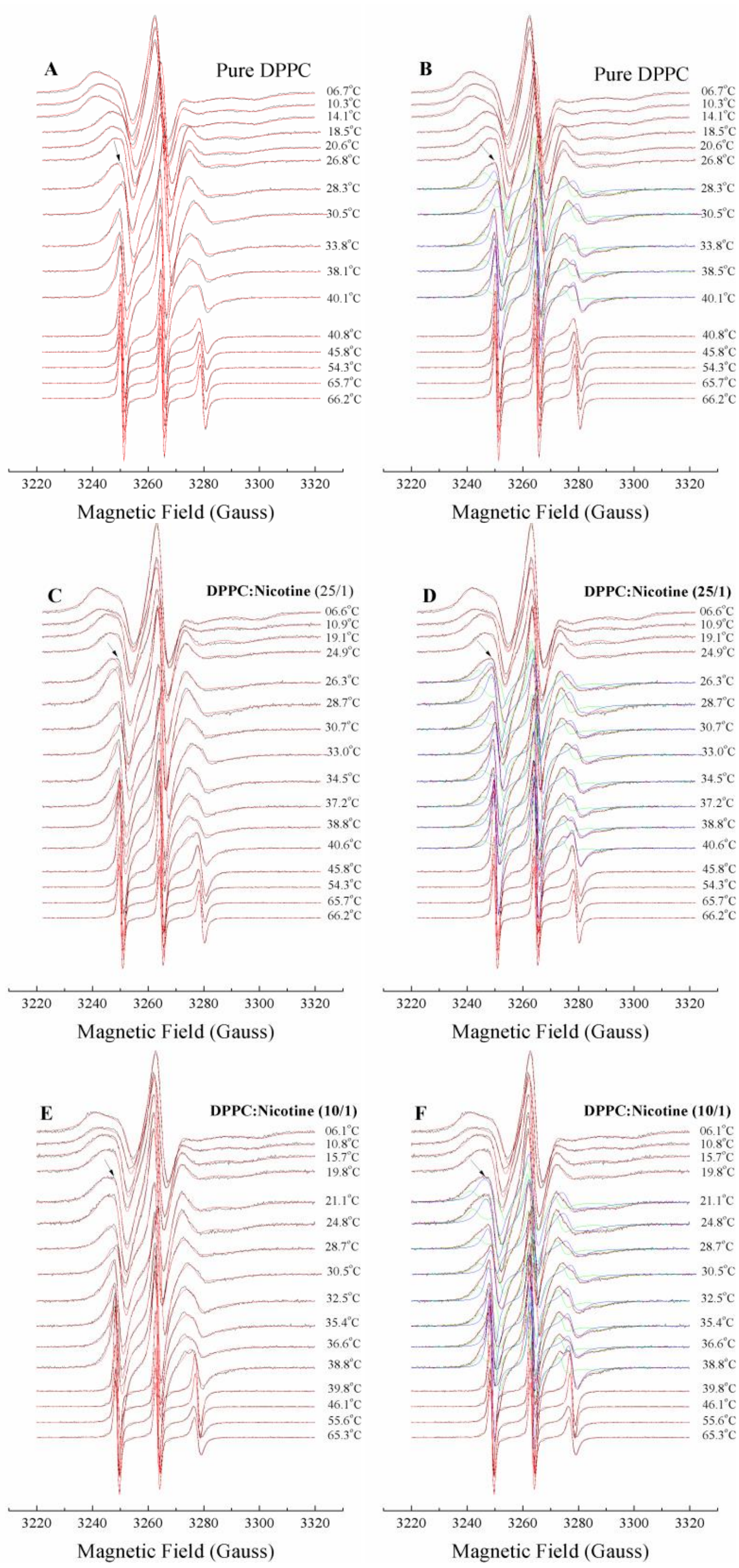

**Table S1.** Activation energies ($E_a$) for the 16-PCSL around an axis perpendicular to the long lipid axis in DPPC and DPPC/nicotine multilamellar vesicles in the gel and fluid phases. The lipid-to-drug molar ratios were 25:1 and 10:1. $E_a$ was calculated from the Arrhenius-type equation $R_{prp}(T) = R_{prp}^0 \times exp(-E_a/RT)$, where $R$ is the gas constant and $T$ is the absolute temperature.

| **Sample** | $E_a^{gel}(kcal/mol/K)$ | $E_a^{fluid}(kcal/mol/K)$ |
|---|---|---|
| blank | 3.5 (0.2) | 8.7 (0.1) |
| 25:1 | 3.6 (0.3) | 8.7 (0.2) |
| 10:1 | 3.6 (0.3) | 8.9 (0.2) |

**Table S2.** Thermodynamic parameters associated with the 16-PCSL probe partitioned between the gel and fluid phases of pure DPPC and DPPC/nicotine vesicles at 25:1 and 10:1 molar ratios. The parameters were obtained by fitting the equation 5 to the experimental data up to the third order for pure DPPC and to the second order for nicotine-containing vesicles.

| **Sample** | **a ($\times 10^2$)** | **b $\times 10^5$ (K)** | **c $\times 10^7$ ($K^2$)** | **d $\times 10^{12}$ ($K^3$)** |
|---|---|---|---|---|
| blank | 570 (90) | -530 (80) | 1600 (300) | -1.7 (0.3) |
| 25:1 | 8.4 (0.6) | -5.1 (0.4) | 7.9 (0.6) | – |
| 10:1 | 5.8 (0.4) | -3.5 (0.3) | 5.4 (0.4) | – |
